# Integrative analysis of Dupuytren’s disease identifies novel risk locus and reveals a shared genetic etiology with BMI

**DOI:** 10.1101/499368

**Authors:** Megan Major, Malika K Freund, Kathryn S Burch, Nicholas Mancuso, Michael Ng, Dominic Furniss, Bogdan Pasaniuc, Roel Ophoff

**Affiliations:** Bioinformatics Interdepartmental Program, University of California Los Angeles, Los Angeles, California, 90095, USA; Department of Human Genetics, David Geffen School of Medicine, University of California Los Angeles, Los Angeles, California, 90095, USA; Department of Pathology and Laboratory Medicine, David Geffen School of Medicine, University of California Los Angeles, Los Angeles, California, 90095, USA; Nuffield Department of Orthopaedics, Rheumatology, and Musculoskeletal Science, University of Oxford, Oxford, OX3 7HE, UK; Department of Plastic and Reconstructive Surgery, Oxford University Hospitals NHS Foundation Trust, John Radcliffe Hospital, Oxford, OX3 9DU, UK; NIHR Biomedical Research Centre, NDORMS, University of Oxford, Oxford, OX3 7HE, UK; Department of Biomathematics, David Geffen School of Medicine, University of California Los Angeles, Los Angeles, California, 90095, USA; Center for Neurobehavioral Genetics, University of California Los Angeles, Los Angeles, California, 90095, USA

**Author notes:** These authors jointly supervised this work (Bogdan Pasaniuc, Roel Ophoff).

## Abstract

Dupuytren’s disease is the common inherited tissue-specific fibrotic disorder. It’s characterized by progressive and irreversible fibroblastic proliferation affecting the palmar fascia of the hand, with an onset typically in the sixth decade of life. Although genome-wide association studies (GWAS) have identified 24 genomic regions associated with Dupuytren’s risk, the biological mechanisms driving signal at these regions remain elusive. We identify potential biological mechanisms for Dupuytren’s disease by integrating the most recent, largest GWAS (*n_cases_* = 3,871, *n_controls_* = 4,686) with eQTLs (47 tissue panels from five consortia, total *n* = 3,975) to perform a transcriptome-wide association study (TWAS). We identify 43 tissue-specific gene associations with Dupuytren’s risk, one of which resides at least 0.5 Mb away from the 24 risk regions previously identified. We also estimate the genome-wide genetic correlation between Dupuytren’s disease and 45 complex traits and find significant genetic correlations between Dupuytren’s disease and body mass index (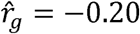, *P* = 1.6 × 10^−6^), type II diabetes (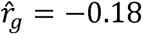, *P* = 1.7 × 10^−4^), triglycerides (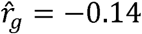, *P* = 3.5 × 10^−4^), and high-density lipoprotein (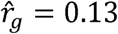, *P* = 4.1 × 10^−4^), which suggests a shared genetic etiology. We further refine the genome-wide genetic correlation signal to identify 8 regions significantly negatively correlated with BMI and 3 regions significantly correlated (1 positively and 2 negatively correlated) with HDL; none of these regions contained the novel gene association identified by TWAS. Our results are consistent with previous epidemiological findings which show that lower BMI increases risk for Dupuytren’s disease. These 12 novel risk regions provide new insight into the biological mechanisms of Dupuytren’s disease and serve as a starting point for functional validation.

## Introduction

Dupuytren’s disease (DD [MIM: 126900]) is a common and disabling connective tissue disorder affecting 5-25% of individuals of European ancestry, characterized by progressive and irreversible fibroblastic proliferation affecting the palmar fascia of the hand^1,2^. DD initially manifests as nodules in the palm of the hand, resulting in contraction and ultimately flexion contractures of the digits in a proportion of individuals affected with DD. Recent twin studies estimate the heritability (i.e., proportion of phenotypic variation explained by genetics) of DD to be ~80%^3^. The largest previous genome-wide association study (GWAS) of DD in individuals of European ancestry identified 26 genome-wide significant single-nucleotide polymorphism (SNP) associations in 24 independent risk regions^4^, and estimated the proportion of phenotypic variance attributable to additive effects of common variants (i.e., SNP-heritability) to be 0.53^4,5^. The vast majority (23 of 24) of DD associations lie in non-coding genomic regions with only one located in an intron^4^, thus the biological implications of these associations are not immediately clear. Investigations into the mechanisms behind the strongest GWAS association, rs16879765 (*P_GWAS_* = 7.2 × 10^−41^), located in the intron of *EPDR1*, revealed an effect on expression and protein secretion of the nearby gene *SFRP4*^4^ but implicated *EPDR1* functionally^6^. Overall, the regulatory mechanisms driving signal at the GWAS associations on DD remains unknown.

In this study, we aimed to explore genetic mechanisms at known risk regions for DD, identify complex traits with possible shared genetic etiologies, and find novel risk regions for DD. Recently, transcriptome-wide association studies^7,8^ (TWAS) have emerged as a way to identify associations between gene expression and a trait. We performed a multi-tissue TWAS by combining a recent DD GWAS^4^ with expression quantitative trait loci^9–14^ (eQTL), integrating gene expression from five consortia in 43 unique tissues, to test for association between expression and DD in 15,198 genes. We identified 43 associations between tissue-specific gene expression and DD, including one novel risk region on chromosome 17. Next, we aimed to understand the genetic relationship between DD and 45 other complex phenotypes by identifying traits that have genetic correlation (i.e., the similarity in genetic effects across two traits) providing etiological insights and plausible causal relationships to investigate^15–18^. We performed genetic correlation analyses through cross-trait linkage disequilibrium (LD) score regression^15^ (LDSC), and find that body mass index (BMI), type II diabetes (T2D), triglycerides (TG), and high-density lipoprotein (HDL) levels are significantly genetically correlated with DD. Additionally, we sought to further refine and understand these relationships with DD and BMI, T2D, TG, and HDL by exploring local regions with enrichments of genetic correlation using ρ-HESS^16^, and found 8 risk regions significantly correlated with BMI and 3 risk regions significantly correlated with HDL. Finally, we aimed to identify a tissue or cell type to prioritize when studying DD.

## Materials and Methods

### DD GWAS summary statistics

Results from a GWAS of DD in UK Europeans (3,871 cases and 4,686 controls) were previously reported^4^. This GWAS summary data contained association statistics for 7,218,238 SNPs, with 6,991,033 SNPs that were imputed from individuals of European ancestry in the Haplotype Reference Consortium^19^. We excluded multi-allelic SNPs, SNPs with ambiguous alleles (e.g., A to T or C to G), and SNPs without an rsID defined by dbSNP144, resulting in 6,126,071 SNPs for downstream analyses.

We used PLINK^20^ to compute independent risk regions (at least one SNP with *P_GWAS_* ≤ 5 × 10^−8^) in the DD GWAS data by clumping SNPs into regions based on LD and distance, using *R^2^* thresholds of 0.3 and 0.25 for between-block LD and within-block LD, respectively. This resulted in 24 independent risk regions.

### TWAS reference panels and details

To find novel risk genes and biologically meaningful associations, we performed a TWAS to test genes expression levels for association with DD. We used FUSION^7^ software (see Web Resources) along with prepackaged gene expression weights. Briefly, TWAS identifies candidate risk genes for DD by integrating results from GWAS and reference panels of gene expression measurements from eQTL studies to associate cis-regulated expression with DD, while accounting for LD. Weights for gene expression were from the Genotype-Tissue Expression Project^9^ v6 (GTEx; 43 tissues, *n* = 449), the Metabolic Syndrome in Men study^10^ (METSIM; adipose, *n* = 563), the Young Finns Study^11,12^ (YFS; blood, *n* = 1,264), the CommonMind Consortium^13^ (CMC; dorsolateral prefrontal cortex, *n* = 452), and the Netherlands Twin Registry^14^ (NTR; blood, *n* = 1,247) reference panels. This totaled to 47 different reference tissue panels that represent 43 unique tissues (see Supplementary Table 1). Description of quality control procedures for these expression data have been previously described^7,21^.

For each reference gene expression panel, FUSION estimates the strength of association between predicted expression of a gene and DD (*Z_TWAS_*) as a function of the vector of GWAS summary Z-scores at a given cis-region (*Z_GWAS_*) and the LD-adjusted weights vector learned from the gene expression data. A p-value (*P_TWAS_*) is obtained using a two-tailed test under *N*(0,1). This process was repeated for each reference tissue panel and gene, resulting in 98,147 tissue-specific gene models involving 15,189 genes (see Supplementary Table 1). We assessed significance with the family-wise error rate threshold at 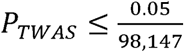.

### Genome-wide genetic correlation with cross-trait LDSC

We estimate genome-wide genetic correlation between DD and 45 complex traits to identify shared genetic risk for DD with other complex traits. To this end, we used cross-trait LDSC^15^, a method for estimating genome-wide genetic correlation between two traits that requires only GWAS summary statistics and reference panel LD (European ancestry from the 1000 Genomes (1000G) Project^22^). We defined the genetic correlation 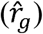 between DD and another trait as significant if it passed the Bonferroni-corrected threshold of 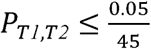.

### Local genetic correlation and putative causality using ρ-HESS

For each of the traits with significant genome-wide genetic correlation with DD, we run ρ-HESS^16^ to estimate the local genetic correlation between each trait and DD within 1702 approximately independent regions genome-wide^23^. For reference LD, we used European ancestry from the 1000G Project^22^. The local genetic correlation 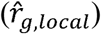 between two traits at a given region was defined as significant if it passed the threshold of 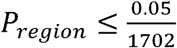.

We also aimed to find evidence for putative causal relationships between DD and other genetically correlated traits. We used the implementation in ρ-HESS based on a previously described method^18^ to prioritize putative causal models between pairs of complex traits. Essentially, for two complex traits, the local genetic correlation is evaluated at regions harboring genome-wide significant GWAS signals specific to each trait. The local genetic correlation for all trait 1 specific regions are summed 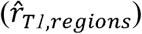 and the local genetic correlation for all trait 2 specific regions are summed 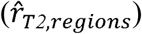. Confidence intervals are determined by 1.96 times jackknife standard error on each side; significance is determined if the confidence intervals do not overlap. The intuition behind this test is that if trait 1 causally influences trait 2 then trait 1 specific regions would have strong genetic correlation with trait 2 but trait 2 specific regions would not have strong genetic correlation with trait 1. Thus, we can leverage the difference in correlations for trait-specific signal at these regions to see if the correlations are consistent with a suggestive causal model^16,18^.

### Tissue and cell type prioritization

To identify tissues and/or cell types that are biologically relevant to DD, we used stratified LD score regression to estimate the enrichment of DD SNP-heritability in 205 publicly available specifically expressed gene (SEG) annotations, each of which represents a set of genes that are specifically expressed in a single tissue or cell type (LDSC-SEG)^24^. Briefly, the 205 annotations were originally created from two datasets: RNA-seq gene expression measurements in 53 human tissues from GTEx v6p^9^ (average of 161 samples per tissue), and a microarray gene expression dataset comprised of 152 tissues and cell types from either human, mouse, or rat (the “Franke Lab” dataset)^25,26^. For each set of specifically expressed genes, an annotation was created by adding 100-kb windows upstream and downstream from the transcribed region of each gene. In addition, we tested for enrichment of DD SNP-heritability in a set of 489 publicly available tissue- or cell type-specific chromatin annotations^24^. 396 of these annotations were originally created from five activating histone marks (H3K27ac, H3K4me3, H3K4me1, H3K9ac, and H3K36me3) and DNase I hypersensitivity (DHS) regions that were present in a subset of 88 tissues and cell types in the Roadmap Epigenomics Consortium^27^. An additional 93 annotations were created from a set of four activating histone marks (H3K27ac, H3K4me3, H3K4me1, and H3K36me3) in 27 tissues from EN-TEx^28^ that were also present in GTEx. Details on the construction of both the SEG annotations and chromatin-based annotations can be found in the original study^24^. Each annotation was tested individually for enrichment of DD SNP-heritability on top of the baseline-LD model^29^ by assessing whether the expected additional per-SNP heritability contribution due to the annotation is significantly nonzero (FDR < 0.1).

We also employed the web application FUMA^30^ in the aim of finding tissues or cell types with differentially expressed genes relevant to DD. FUMA maps GWAS results to create a gene set in three ways: (1) physical proximity on the genome, (2) eQTL associations, and (3) chromatin interaction. We used the gene property analyses (implemented from MAGMA^31^) and differentially expressed gene (DEG) analysis to prioritize different tissues or cell types. For the gene property analysis, FUMA tests if expression of the GWAS gene set in a single tissue or cell type is statistically different than the average expression of the GWAS gene set across all tissues or cell types. We perform this gene property analysis in 53 GTEx^9^ tissues 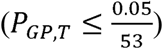 as well as in 5115 study-defined cell types 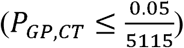 using single cell RNA-seq data from 28 studies^32–58^ as described on the FUMA website (see Web Resources). For the DEG analysis, FUMA defines differentially expressed genes in each tissue by performing a two-sided t-test for that one tissue against all other tissues. Each of the 53 GTEx^9^ tissues is tested for up-regulation, down-regulation, and both-sided DEG sets. We removed tissues where DEG sets had less than 30 genes to avoid underpowered correlations; significance was defined by Bonferroni correction for the number of tests 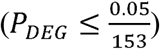.

## Results

### TWAS identifies 18 risk genes for DD

To explore putative biological mechanisms at known DD risk regions, we performed a multi-tissue TWAS to identify genes (specifically, cis-regulated gene expression), associated with DD (see Materials and Methods). Briefly, TWAS identifies candidate risk genes for DD by integrating results from GWAS and reference panels of gene expression measurements from eQTL studies to associate cis-regulated expression with DD, while accounting for LD. We used tissue reference panels from GTEx^9^, METSIM^10^, YFS^11,12^, CMC^13^, and NTR^14^ resulting in 47 different reference tissue panels with a combined sample size of 3,975 (see Materials and Methods; Supplementary Table 1). Using these reference panels, we tested 98,147 tissue-specific gene models and found 43 significant tissue-specific gene-trait associations at a Bonferroni-corrected threshold of 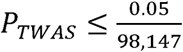 (Table 1, Supplementary Table 2). GWAS SNP association strength and TWAS tissue-specific gene model association strength can be seen in Figure 1. These 43 significant models were composed of 18 genes among 23 tissue panels–7 genes were significant in multiple tissues (Table 1). 36 of the 43 significant tissue-specific gene models were within 0.5Mb of any of the previously identified 24 risk regions.

**Figure 1:**
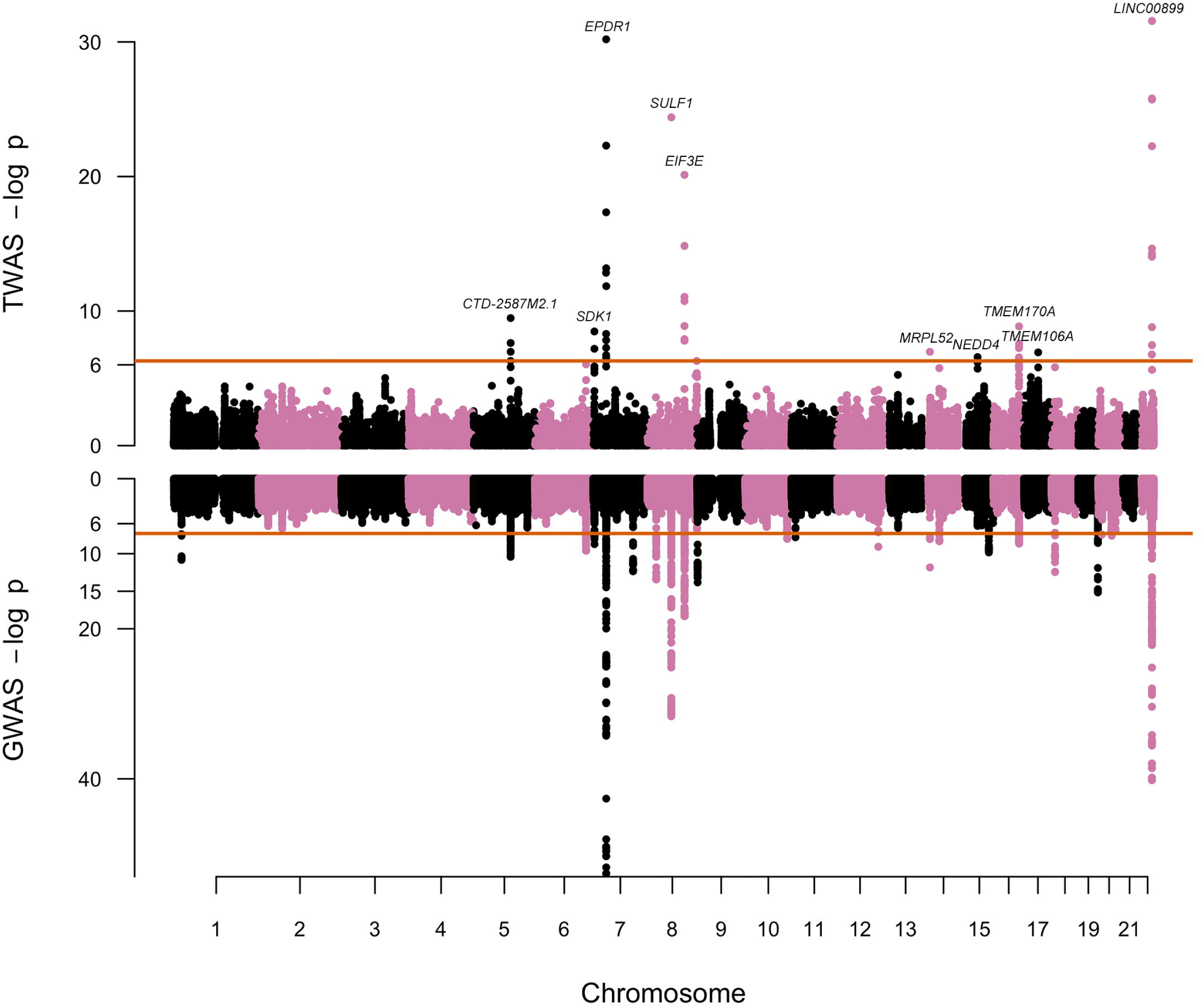
DD TWAS and GWAS associations. Shown here are Manhattan plots for TWAS associations (top) and GWAS associations (bottom). For TWAS associations, each point corresponds to an association test between tissue-specific predicted gene expression and DD, with the orange line representing the threshold for significance in log-scale (*P_TWAS_* ≤ 5.09 × 10^−7^). The most significant tissue-specific gene model for each peak is labeled by gene. For GWAS associations, each point corresponds to an association test between a SNP and DD, with the orange line representing the traditional genome-wide significance threshold in log-scale (*P_GWAS_* ≤ 5 × 10^−8^).

**Table 1:**
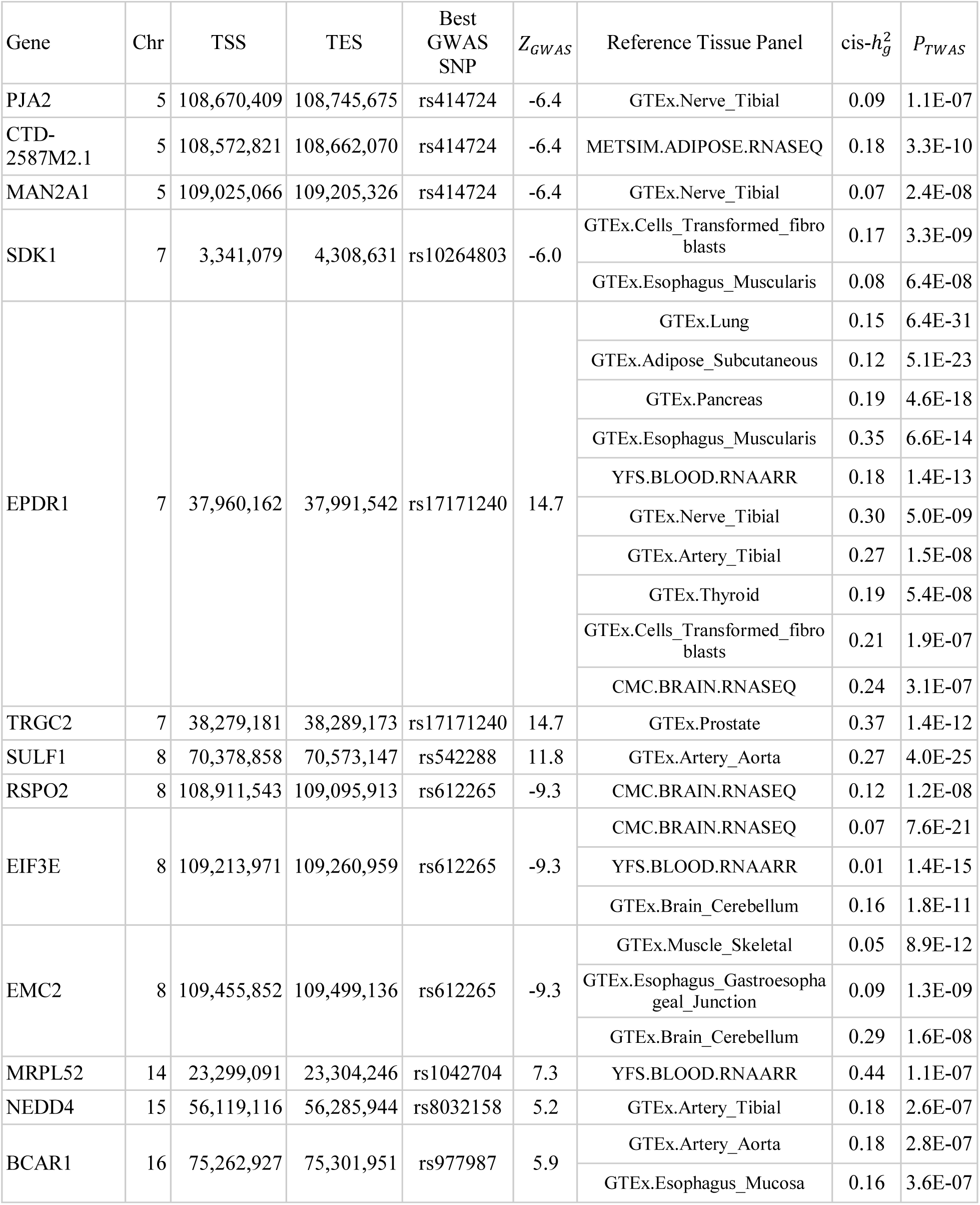
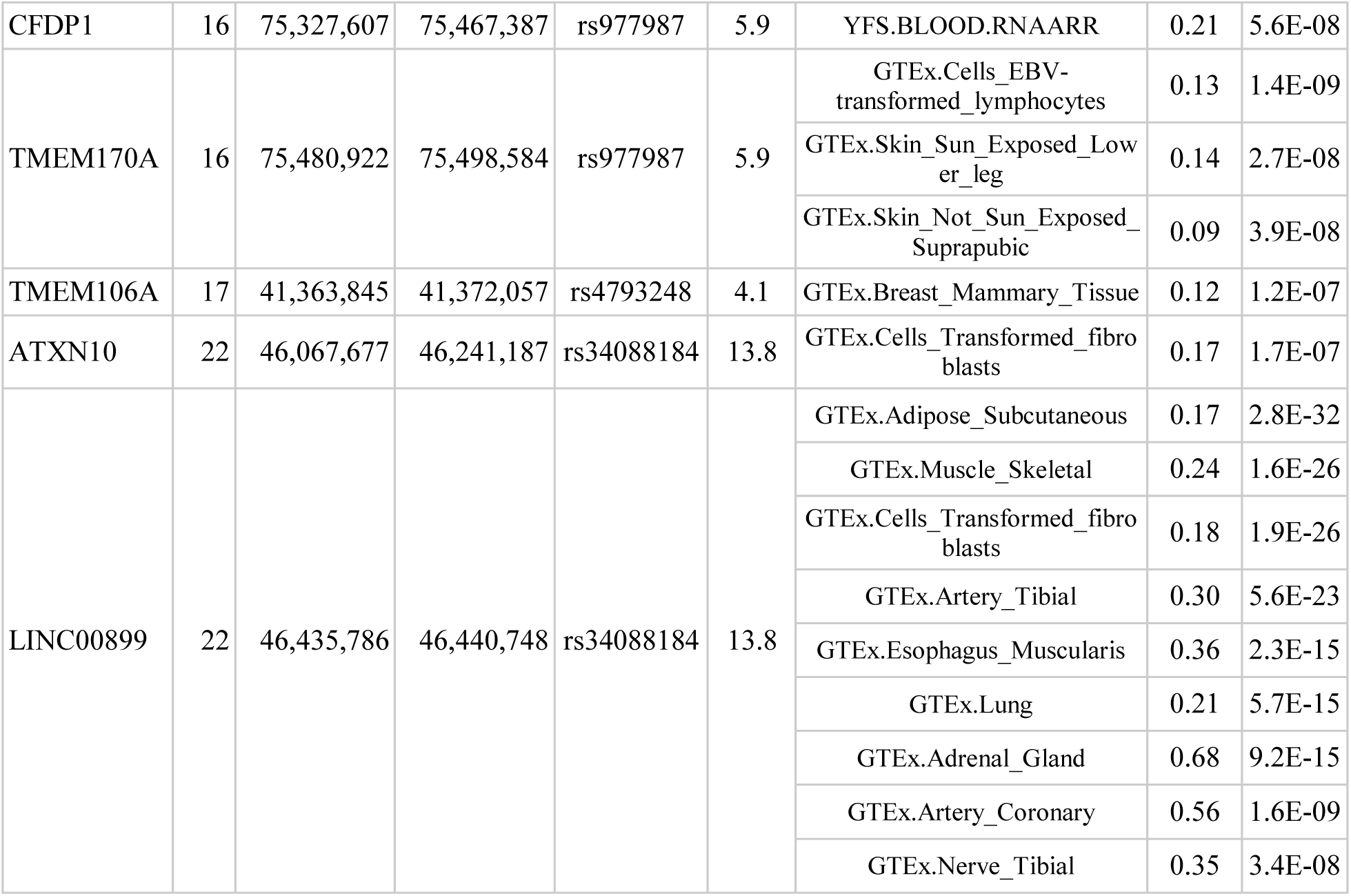
43 significant tissue-specific gene expression models from TWAS. These are the 43 significant (*P_TWAS_* ≤ 0.05/98,147) tissue-specific gene models across 18 genes.

One region of interest is on chromosome 7, where *EPDR1* was found to be significant in 10 different tissue panels (most significant in lung tissue, *P_TWAS_* = 6.4 × 10^−31^). This region has been previously investigated because of its strong association signal (Odds Ratio 1.93 and *P_GWAS_* = 7.2 × 10^−41^) with DD^4^. The variant with the strongest association in this region, rs16879765, is in an intron of *EPDR1*. Although decreased secretion of the nearby WNT-agonist *SFRP4* was correlated with the high risk genotype^4^, genetic and functional evidence point toward *EPDR1* being the disease-relevant gene for this region, which has been functionally validated as contributing to myofibroblast contractility^6^. All three transcripts of *EPDR1* are found in affected DD tissue and knockdown of *EPDR1* attenuates contractility in fibroblast-populated collagen lattice assays^6^.

### TWAS identifies novel risk region on chromosome 17

To identify possible novel risk regions from TWAS associations, we aimed to see if any tissue-specific gene models were independent of established GWAS associations. After grouping association signal into 1Mb regions, we found 13 regions with only significant GWAS SNP(s), 1 region with only significant TWAS model(s), and 11 regions with both significant GWAS SNP(s) and TWAS model(s). Here we define a region identified through TWAS to be novel if (1) the strongest DD associated SNP in the gene’s region is not genome-wide significant (i.e., *P_GWAS_* ≥ 5 × 10^−8^) and (2) that the TSS of the TWAS-gene is not within 0.5Mb of the previously known 24 risk regions. With these constraints, we identified one novel risk region for DD (Figure 2). To ensure our result was robust to long-range LD, we expanded our window criteria to include 1Mb and 2Mb and found no change. We found a single tissue-specific gene model, *TMEM106A* (*P_TWAS_* = 1.2 × 10^−7^; GTEx breast mammary tissue), was significantly associated with DD risk at this region (Figure 2). To determine that the *TMEM106A* association was robust to possible LD confounding, we performed a permutation test using GWAS summary statistics and found similar results (*P_perm_* = 8.91 × 10^−3^). There were 12 tissue panels that expression for *TMEM106A* was modeled from (Supplementary Table 3).

**Figure 2:**
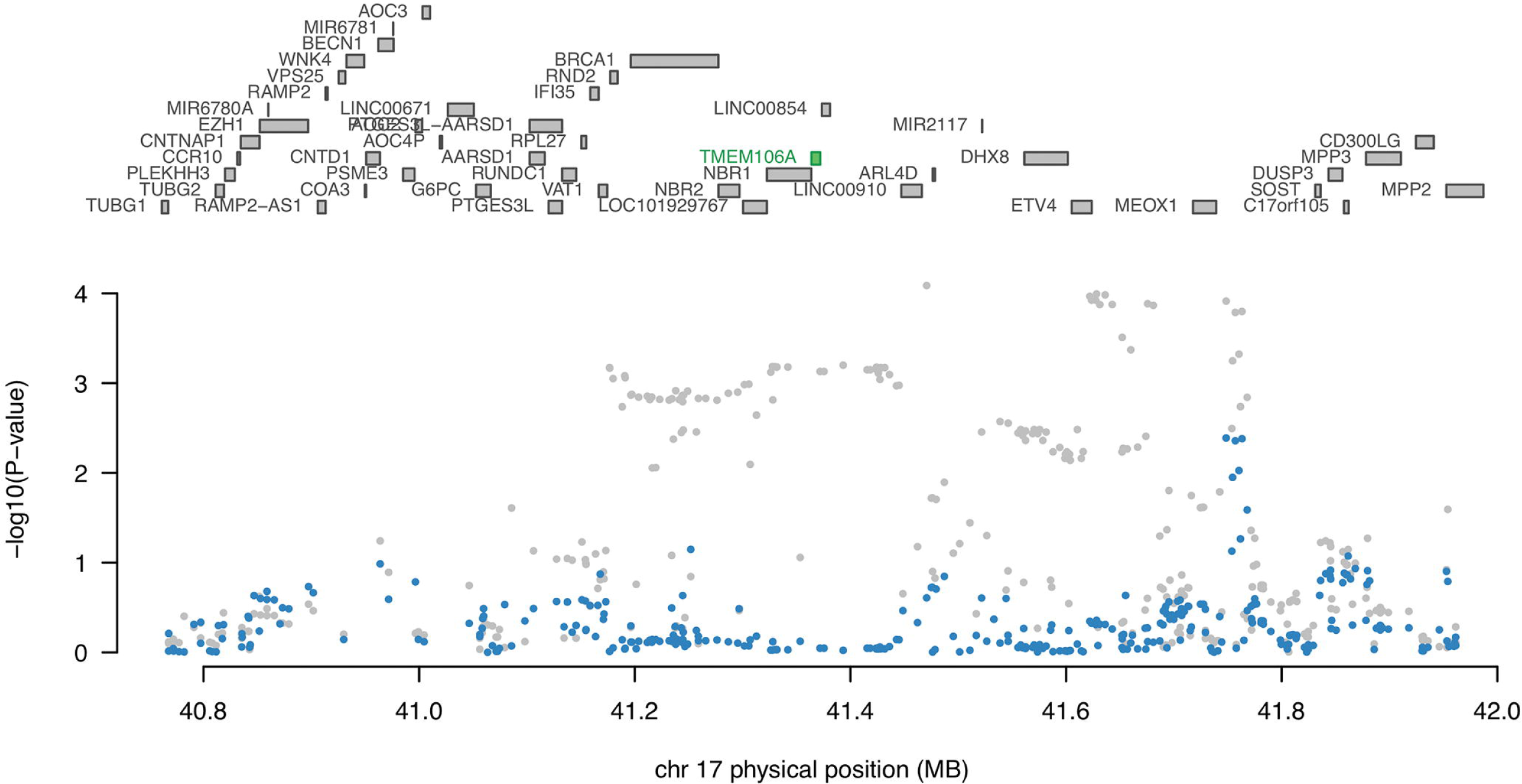
Novel risk region identified on chromosome 17. Shown here is the novel risk region identified through TWAS; the grey points are GWAS SNPs association strength and the blue points are the GWAS SNPs association strength conditioned on the *TMEM106A* expression model (green, significant in GTEx breast mammary tissue). This tissue-specific model was still significant under 1,347 permutations (*P_perm_* = 8.9 × 10^−3^).

### Estimates of SNP-heritability in DD are higher than previously proposed

We obtained a SNP-heritability estimate of 0.67 (s.e. = 0.08), using LDSC^59^. We also used Heritability Estimator from Summary Statistics (HESS), a previously described method using similar framework as ρ-HESS that estimates local SNP-heritability, and found the total SNP-heritability to be 0.532 (s.e. = 0.282), similar to the previous estimate of 0.533 using GCTA^4,5^. Because HESS is optimized for GWAS with sample sizes greater than 50,000 (contributing the large standard error), we included the LDSC regression estimate of SNP-heritability when running HESS to obtain more stable estimates of local SNP-heritability^59,60^.

### Genetic correlation suggests shared genetic etiology with DD

To identify traits that have a shared genetic etiology with DD, we used cross-trait LDSC^15^ which estimates the genetic correlation between two traits using GWAS summary statistics (see Materials and Methods). Results for the genetic correlation test between DD and 45 other traits (average sample size 132,115) can be found in Table 2. These 45 traits include a variety of anthropometric, immune, hematological, neurological, and cardiovascular related traits and disorders. Four traits were found to have significant genetic correlation with DD (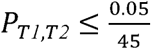, identical results correcting for a FDR < 0.1): body mass index (BMI), 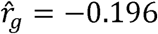; high density lipoprotein (HDL), 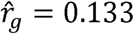; triglycerides (TG), 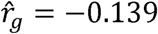; and type II diabetes (T2D), 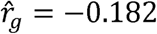 (Table 2). These results are compatible with previous observational studies^2,61,62^. Notably, the negative genetic correlation between BMI and DD is consistent with a previous epidemiological investigation showing that the risk of DD was inversely proportional to BMI, after correcting for age, race, and sex in 14,844 patients diagnosed with DD^63^.

**Table 2:**
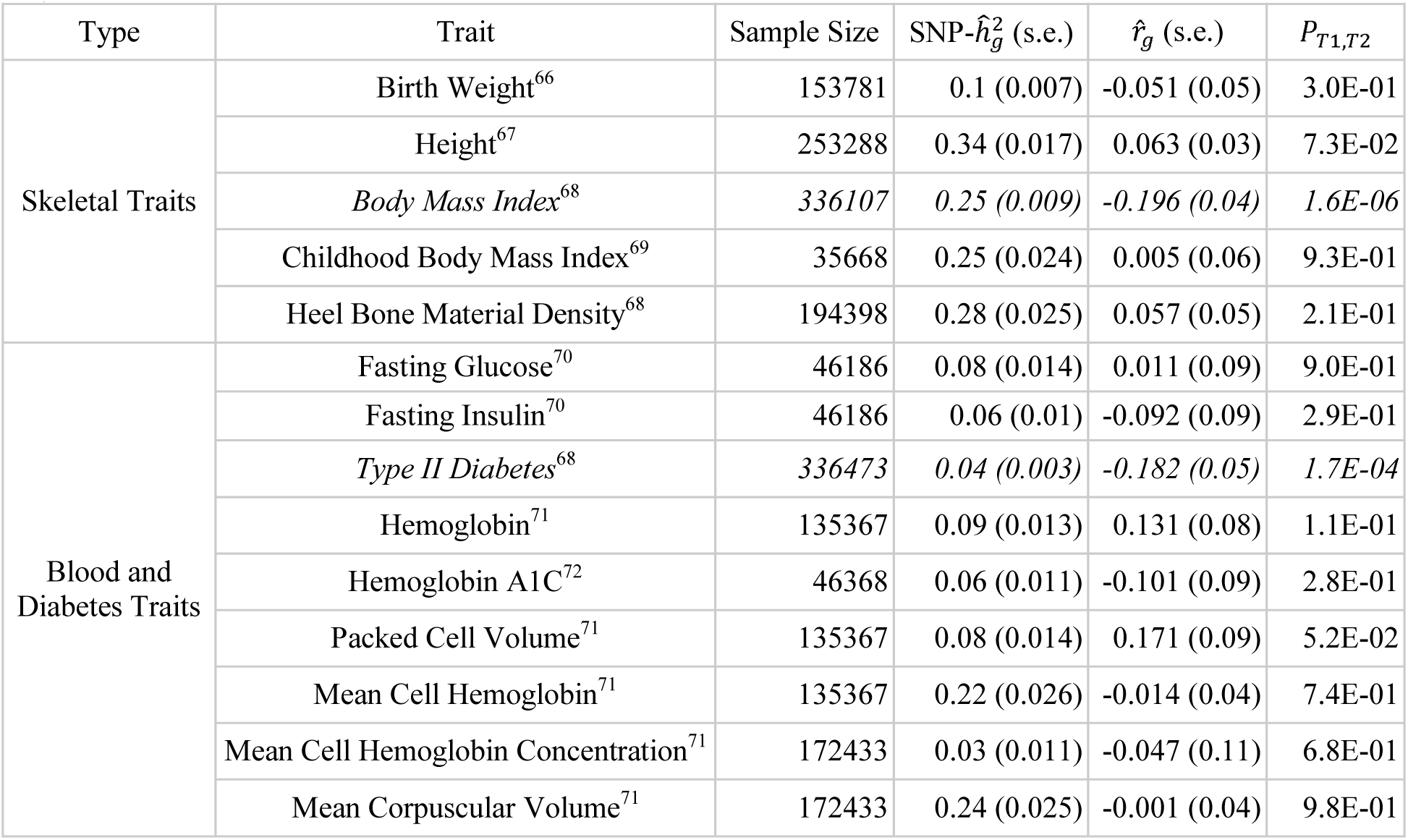
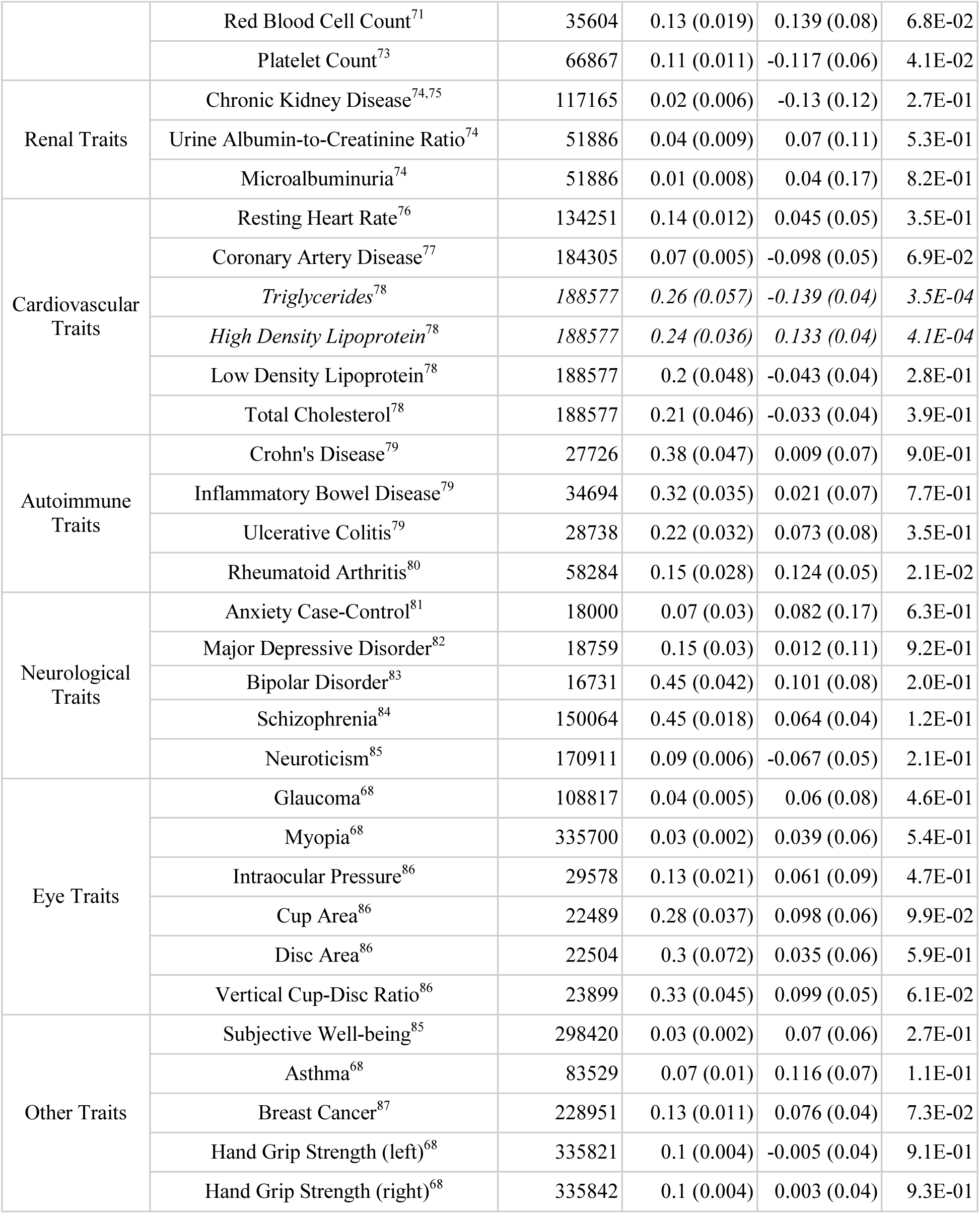
Genetic correlation results between DD and 45 other traits. We have grouped related traits under the “Type” column. The four traits that were significantly (*P_T1,T2_* ≤ 0.05/45) correlated with DD are shown in italics.

### Local genetic correlation analysis yields 11 regions to further study

Having found evidence for BMI, HDL, TG, and T2D sharing genetic factors with DD at a genome-wide scale, we next aimed to locate possible shared genomic regions. We did this by running ρ-HESS^16^ to estimate the local genetic correlation between DD and each of the four traits in 1702 approximately independent LD blocks^23^ (see Materials and Methods). The genome-wide genetic correlation results from cross-trait LDSC and ρ-HESS are fairly consistent (Pearson’s r = 0.94; Supplementary Table 4); differences may have resulted from using metabochip array^64^ GWAS statistics with LDSC (HDL and TG), which is discouraged for cross-trait LDSC^15^, as well as smaller sample size in the DD GWAS adding noise to estimates from ρ-HESS^16,60^. We found eight regions significantly genetically correlated 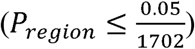 between DD and BMI, three regions between DD and HDL, and no regions between DD and TG or between DD and T2D (Table 3). Of these 11 regions, three contained a genome-wide significant association in the DD GWAS^4^. Only one of the 11 regions contained significant tissue-specific gene models from TWAS; the 10 models for *EPDR1* were within the DD and HDL genetically correlated 7:37555184-38966703 region (Supplemental Table 2, Table 3).

**Table 3:**
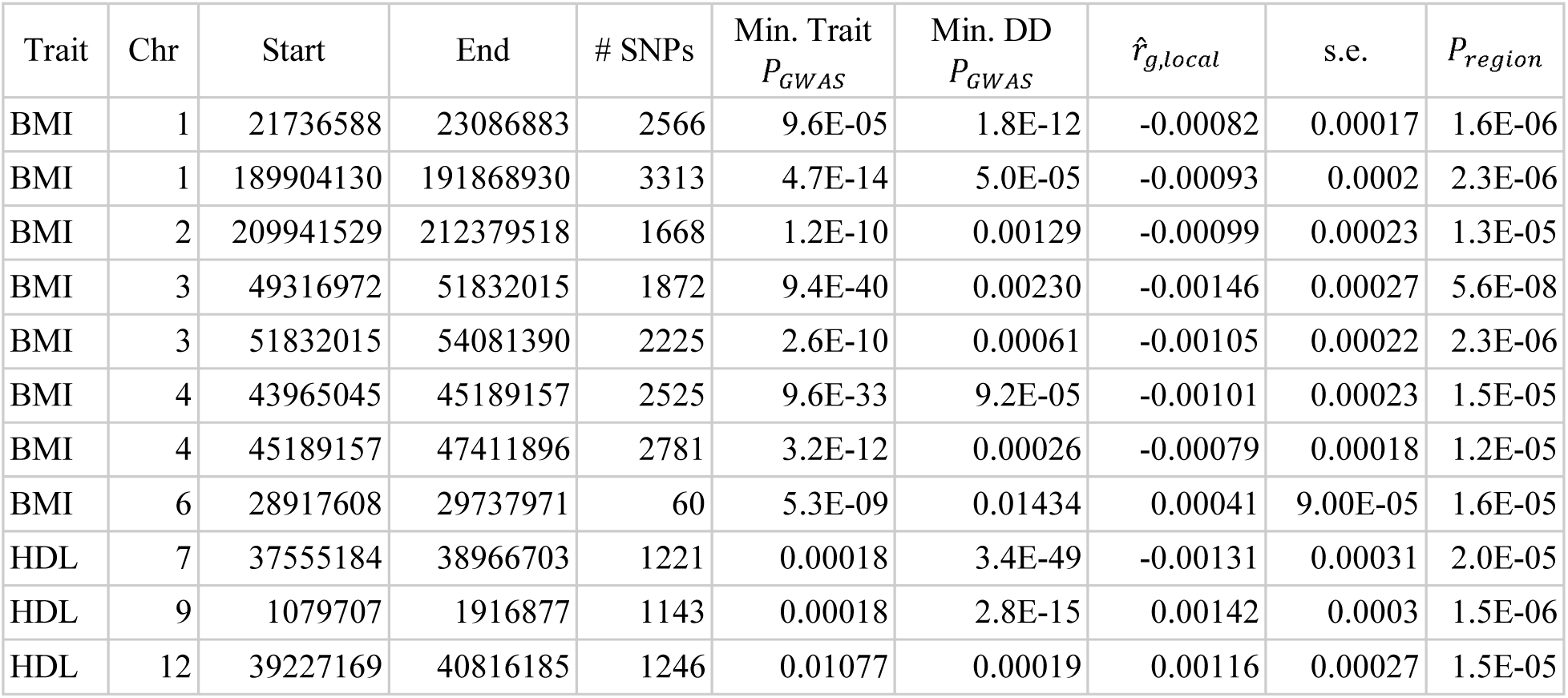
Regions with significant genetic correlation between DD and other traits. This table lists the eight regions demonstrating significant genetic correlation between DD and BMI, and the three regions demonstrating significant correlation between DD and HDL; significance was assessed at a Bonferroni-corrected threshold of *P_region_* ≤ 0.05/1702 for each trait. Also included is the number of SNPs within each region (“# SNPs”) as well as the minimum GWAS association p-value for either BMI or HDL (“Min. Trait *P_GWAS_*”) and DD (“Min. DD *P_GWAS_*”). All other regions demonstrated no significant genetic correlation between DD and any trait tested.

| Trait | Chr | Start | End | # SNPs | Min. Trait<br>$P_{GWAS}$ | Min. DD<br>$P_{GWAS}$ | $\hat{r}_{g,local}$ | s.e. | $P_{region}$ |
| --- | --- | --- | --- | --- | --- | --- | --- | --- | --- |
| BMI | 1 | 21736588 | 23086883 | 2566 | 9.6E-05 | 1.8E-12 | -0.00082 | 0.00017 | 1.6E-06 |
| BMI | 1 | 189904130 | 191868930 | 3313 | 4.7E-14 | 5.0E-05 | -0.00093 | 0.0002 | 2.3E-06 |
| BMI | 2 | 209941529 | 212379518 | 1668 | 1.2E-10 | 0.00129 | -0.00099 | 0.00023 | 1.3E-05 |
| BMI | 3 | 49316972 | 51832015 | 1872 | 9.4E-40 | 0.00230 | -0.00146 | 0.00027 | 5.6E-08 |
| BMI | 3 | 51832015 | 54081390 | 2225 | 2.6E-10 | 0.00061 | -0.00105 | 0.00022 | 2.3E-06 |
| BMI | 4 | 43965045 | 45189157 | 2525 | 9.6E-33 | 9.2E-05 | -0.00101 | 0.00023 | 1.5E-05 |
| BMI | 4 | 45189157 | 47411896 | 2781 | 3.2E-12 | 0.00026 | -0.00079 | 0.00018 | 1.2E-05 |
| BMI | 6 | 28917608 | 29737971 | 60 | 5.3E-09 | 0.01434 | 0.00041 | 9.00E-05 | 1.6E-05 |
| HDL | 7 | 37555184 | 38966703 | 1221 | 0.00018 | 3.4E-49 | -0.00131 | 0.00031 | 2.0E-05 |
| HDL | 9 | 1079707 | 1916877 | 1143 | 0.00018 | 2.8E-15 | 0.00142 | 0.0003 | 1.5E-06 |
| HDL | 12 | 39227169 | 40816185 | 1246 | 0.01077 | 0.00019 | 0.00116 | 0.00027 | 1.5E-05 |

### Genetic correlation patterns of BMI/TG and DD consistent with putative causality

To further elucidate the relationships of these traits with DD, we used ρ-HESS^16^ to test for evidence of putative causality through GWAS estimated genetic effects for BMI, HDL, TG, and T2D acting on DD or vice versa (see Materials and Methods). Both BMI and TG showed suggestive patterns that would be consistent with a putative causal relationship with DD, while HDL and T2D did not (Figure 3, Supplemental Figure 1). For example, when considering BMI and DD, the correlation at 399 BMI-specific regions (−0.27, s.e. = 0.044) is seemingly stronger than the correlation at 19 DD-specific regions (−0.03, s.e. = 0.15), indicating that regions that increase BMI tend to decrease risk of DD; this is consistent with a model where BMI genetic effects decrease risk of DD (Figure 3). The same is true for TG and DD; the correlation at 65 TG-specific regions (−0.3, s.e. = 0.08) is seemingly stronger than the correlation at 22 DD-specific regions (0.018, s.e. = 0.2; Figure 3). Both of these results are not significant (assessed by overlap of confidence intervals, 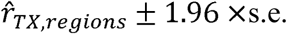); this is most likely because of the relatively reduced sample size in the DD GWAS study. Nonetheless, there is evidence of a putative causal relationship with BMI affecting TG^16,18^ and results from TG and DD may be from a mediated causal relationship of BMI affecting TG, which in turn affects DD (Figure 3).

**Figure 3:**
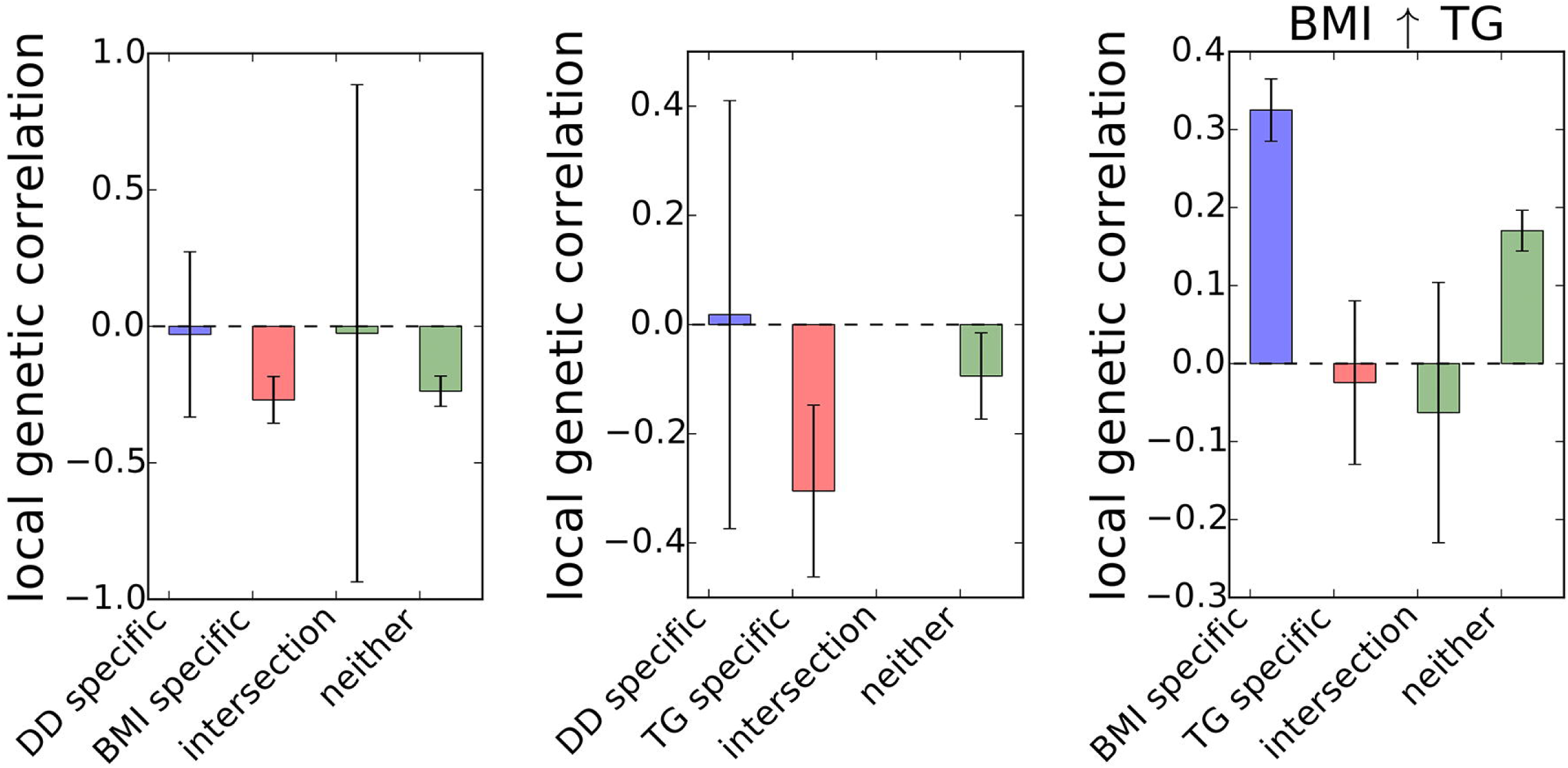
Tentative evidence for putative causality with DD. Here we show the genetic correlation for three different pairs of traits (DD/BMI, left; DD/TG, center; and BMI/TG, right) between four groupings of SNPs: (1) GWAS-significant SNPs specific to trait 1, (2) GWAS-significant SNPs specific to trait 2, (3) GWAS-significant SNPs for both trait 1 and trait 2, and (4) all non-significant SNPs shared between studies. In the left plot, since GWAS-significant SNPs specific to BMI have more enrichment of genetic correlation compared to those specific to DD, we can putatively interpret that BMI SNPs are driving the shared genetic etiology. The same can be said for the middle plot with TG. On the right, for completeness, we show the same correlation for BMI and TG, which was significant. Error bars are defined by the genetic correlation ± 1.96 times the s.e. for each grouping of SNPs.

### DD most relevant tissue or cell type unidentifiable with current data

Finally, we aimed to identify relevant tissues or cell types for DD. First, we used S-LDSC^65^ to estimate the enrichment of SNP-heritability of DD (controlling for the baseline-LD model^29^) in two sets of publicly available annotations^24^: annotations representing specifically expressed genes (SEG) in 205 tissues or cell types^9,25,26^ and 489 annotations representing 6 chromatin features (DHS and 5 histone marks) in 91 tissues or cell types^27,28^. Among the 205 SEG annotations, synovial membrane tissue was the most enriched for DD SNP-heritability on top of the baseline-LD model, but none of the 205 annotations were statistically significant (FDR < 0.1; Supplementary Table 5). Among the 489 chromatin annotations, we found that esophageal-mucosa tissue was the most enriched for DD SNP-heritability, however none of 489 annotations were statistically significant (FDR < 0.1; Supplementary Table 6). Next, we prioritized tissues and cell types using FUMA^30^, a platform to visualize and interpret GWAS summary statistics (see Materials and Methods). After using FUMA to create a gene set from the GWAS statistics, we first performed a gene property analysis (which tests if gene expression in a single tissue or cell type is statistically different than the average gene expression across all tissues or cell types) in 53 tissue types^9^. Although none of the 53 tissues showed a significant effect 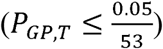, the effect was strongest in cell transformed fibroblasts (Supplementary Table 7). We then assessed whether the GWAS gene set was enriched in any of the differentially expressed gene (DEG) sets for tissues. The up-regulated DEG sets for tibial artery and aorta tissues both demonstrated significant (*P_DEG_* = 5.5 × 10^−5^ and 7.8 × 10^−5^, respectively) overlap with the GWAS gene set (Supplementary Table 8). We also performed a gene property analysis using cell type specific expression data for 5115 study-defined cell types from 28 scRNA-seq studies^32–58^. While none of the single cell types were significant 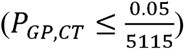, stromal cells and muscle cells were among the top five results (Supplementary Table 9). As a final analysis, we averaged the *χ^2^*-statistic (*Z_TWAS_^2^*) for the 43 significant TWAS models within each tissue to determine which tissue had the most enrichment of TWAS signal. We found adipose subcutaneous tissue was most enriched among the 23 tissues with significant TWAS models (Supplementary Figure 2). Because of the lack of consistency between methods and lack of statistical significance in many methods, we are lead to believe that likely the relevant tissue or cell type is not represented in current datasets.

## Discussion

In this work, we aimed to better understand the genetic architecture of DD, find plausible biological mechanisms at known risk regions for DD, understand the relationship between DD and a variety of other traits, and identify possible novel risk regions through local genetic correlation with other traits or genetic-mediated gene expression effects. We highlight that the estimated SNP-heritability of DD (0.53-0.67) is relatively close to estimates of heritability from twin studies (0.8). We also note that the strong concentration of DD GWAS signal in a handful of genomic regions is more consistent with an oligogenic architecture than a polygenic one, suggesting that further functional studies could be particularly fruitful as compared to more polygenic traits and diseases. We also identify a negative genetic correlation between DD and BMI, supporting a previous epidemiological study that observationally showed a negative correlation between the traits^63^; understanding the relationship between DD and BMI as well as that between DD and TG could shed light on shared biologically important pathways. Finally, we identify one novel risk region from TWAS, and identify 11 regions with significant local genetic correlation between DD and BMI or HDL. Overall, our findings highlight the need for more investigation into these regions as a first step.

Additionally, we note a few caveats in our results. First, though the sample size of 8,557 for the DD GWAS is the largest yet, it is possible that additional GWAS regions remain undiscovered due to the limit in power and this also would further reduce power to fully detect associations and relationships with other traits. Second, while the patterns of genetic correlation between BMI and DD as well as TG and DD are somewhat consistent with causal relationships, true causality between these traits cannot be determined without functional experimentation. Third, we emphasize that TWAS may not detect the true mechanism of disease if the gene expression is not mediated through genetics or if disease-relevant tissue is not well-represented in available gene expression reference panels. This may be further illustrated by the fact we were unable to identify a specific tissue or cell type to prioritize for further study in DD. This could also be due to the small sample size of the DD GWAS, the cell-type specificity of enhancer elements, or again the publication bias away from musculoskeletal connective tissues, leading to a gap in the available datasets.

Future work should be taken in multiple directions. First, we provide additional evidence that *EPDR1* may contribute to the pathogenesis of DD; further work should be dedicated to functionally validate and understand this gene in connection with DD, as it may represent an attractive therapeutic target. Second, there is strong evidence for a relationship between BMI/TG and DD–elucidating the mechanism may lead to interesting observations with implications for the treatment of both traits. Third, additional GWAS, with larger sample sizes and in additional populations, will uncover more of the contribution of genetic variation to DD. And fourth, given the putative oligogenic architecture of DD, and that our tissue and cell type analyses lacked consistent results, it might also be rewarding to generate more functional -omics data, such as reference gene expression panels or chromatin accessibility data in the palmar fascia tissue. These resources would offer valuable insight into the underlying mechanisms of DD and opportunity to explore therapeutic avenues.

## Supporting information

## Supplemental Data

Supplemental Data includes two figures and nine tables.

## Acknowledgements

This work was supported by National Institutes of Health (NIH) awards R01HG009120, R01MH115676, R01HG006399, and U01CA194393. M.K.F. is funded by NIH-NCI National Cancer Institute T32LM012424; K.S.B. is funded by NIH Training Grant in Genomic Analysis and Interpretation T32HG002536; D.F. is funded by an Intermediate Fellowship from the Wellcome Trust (097152/Z/11/Z). This work was also supported by the NIHR Biomedical Research Centre, Oxford.

## Web Resources

OMIM, https://www.omim.org/
1000 Genomes, http://www.internationalgenome.org/
PLINK, https://www.cog-genomics.org/plink2/
FUSION, http://gusevlab.org/projects/fusion/
GTEx Portal, https://gtexportal.org/home/
LD scores and annotations, https://data.broadinstitute.org/alkesgroup/LDSCORE/
DEPICT, https://data.broadinstitute.org/mpg/depict/depict_download/tissue_expression/
FUMA, http://fuma.ctglab.nl/

