## Supplementary material for "Integrative analysis of Dupuytren’s disease identifies novel risk locus and reveals a shared genetic etiology with BMI"

SUPPLEMENTAL FIGURES

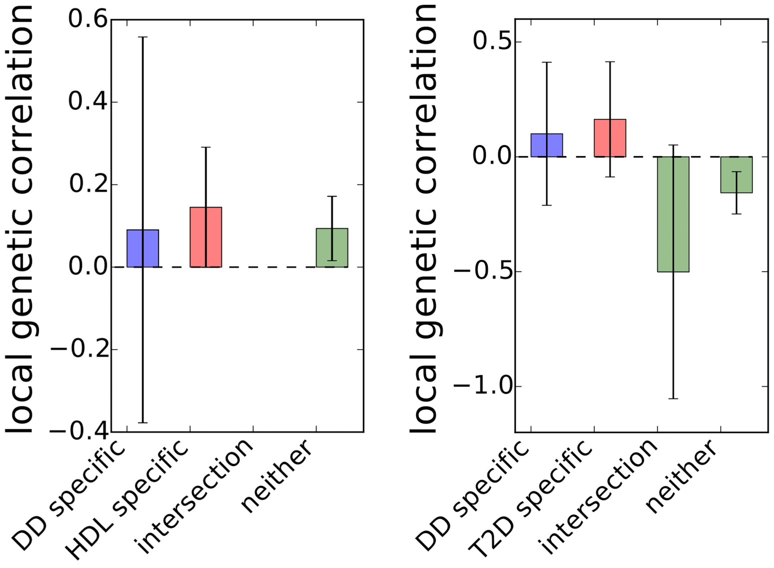


**Figure S1: No evidence for causal relationships of HDL or T2D on DD.** Here we have the genetic correlation for DD and HDL or DD and T2D between four groupings of SNPs: (1) DD specific significant GWAS SNPs, (2) HDL or T2D specific significant GWAS SNPs, (3) significant GWAS SNPs in both DD and HDL or T2D and (4) all non-significant SNPs shared between studies. While there is evidence of some shared genetics from effect of SNPs, the results found here are not consistent with a causal relationship of HDL or T2D on DD. Error bars are defined by the genetic correlation ± 1.96 times the s.e. for each grouping of SNPs.

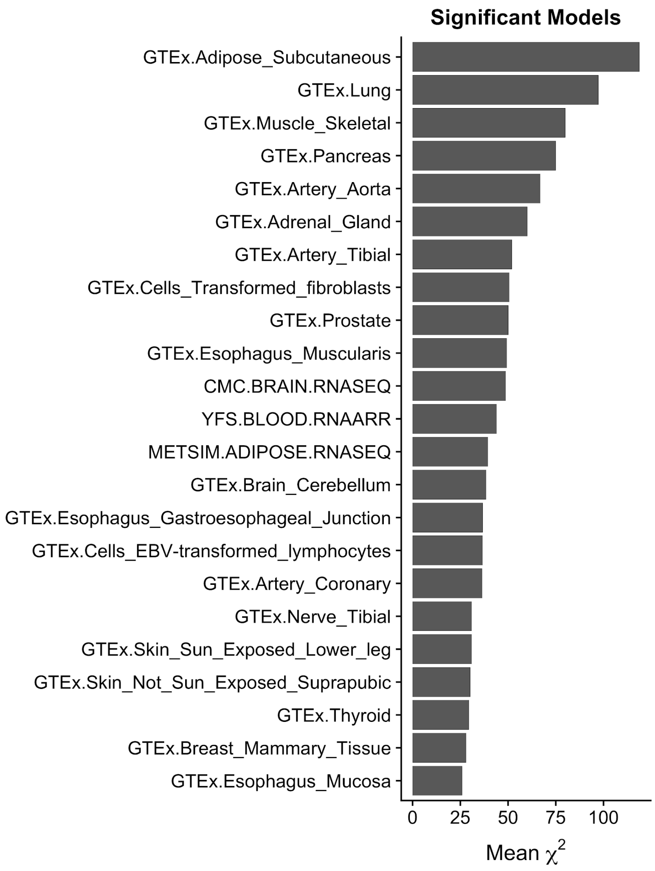


**Figure S2: Mean $\chi^2$ over the 43 significant tissue-specific gene models from TWAS.** To find the tissue with the most signal from TWAS, we calculated the mean $\chi^2$ statistic for all significant models within a tissue.

### SUPPLEMENTAL TABLES

| Reference Panel | Sample Size | Number of Gene Models | Number of Genes with Significant Association |
|---|---|---|---|
| GTEx.Adipose_Subcutaneous | 298 | 3492 | 2 |
| GTEx.Adipose_Visceral_Omentum | 185 | 2037 | 0 |
| GTEx.Adrenal_Gland | 126 | 1457 | 1 |
| GTEx.Artery_Aorta | 197 | 2976 | 2 |
| GTEx.Artery_Coronary | 118 | 1183 | 1 |
| GTEx.Artery_Tibial | 285 | 3760 | 3 |
| GTEx.Brain_Caudate_basal_ganglia | 100 | 921 | 0 |
| GTEx.Brain_Cerebellar_Hemisphere | 89 | 1488 | 0 |
| GTEx.Brain_Cerebellum | 103 | 1982 | 2 |
| GTEx.Brain_Cortex | 96 | 1034 | 0 |
| GTEx.Brain_Frontal_Cortex_BA9 | 92 | 924 | 0 |
| GTEx.Brain_Hippocampus | 81 | 525 | 0 |
| GTEx.Brain_Hypothalamus | 81 | 590 | 0 |
| GTEx.Brain_Nucleus_accumbens_basal_ganglia | 93 | 865 | 0 |
| GTEx.Brain_Putamen_basal_ganglia | 82 | 624 | 0 |
| GTEx.Breast_Mammary_Tissue | 183 | 1974 | 1 |
| GTEx.Cells_EBV-transformed_lymphocytes | 114 | 1420 | 1 |
| GTEx.Cells_Transformed_fibroblasts | 272 | 4198 | 4 |
| GTEx.Colon_Sigmoid | 124 | 1460 | 0 |
| GTEx.Colon_Transverse | 169 | 2064 | 0 |
| GTEx.Esophagus_Gastroesophageal_Junction | 127 | 1446 | 1 |
| GTEx.Esophagus_Mucosa | 241 | 3319 | 1 |
| GTEx.Esophagus_Muscularis | 218 | 3230 | 3 |
| GTEx.Heart_Atrial_Appendage | 159 | 1579 | 0 |
| GTEx.Heart_Left_Ventricle | 190 | 1531 | 0 |
| GTEx.Liver | 97 | 717 | 0 |
| GTEx.Lung | 278 | 2872 | 2 |
| GTEx.Muscle_Skeletal | 361 | 2891 | 2 |
| GTEx.Nerve_Tibial | 256 | 4324 | 4 |
| GTEx.Ovary | 85 | 909 | 0 |
| GTEx.Pancreas | 149 | 1632 | 1 |
| GTEx.Pituitary | 87 | 1115 | 0 |

| | | | |
|---|---|---|---|
| GTEx.Prostate | 87 | 834 | 1 |
| GTEx.Skin_Not_Sun_Exposed_Suprapubic | 196 | 2460 | 1 |
| GTEx.Skin_Sun_Exposed_Lower_leg | 302 | 3636 | 1 |
| GTEx.Small_Intestine_Terminal_Ileum | 77 | 461 | 0 |
| GTEx.Spleen | 89 | 1410 | 0 |
| GTEx.Stomach | 170 | 1455 | 0 |
| GTEx.Testis | 157 | 3155 | 0 |
| GTEx.Thyroid | 278 | 3997 | 1 |
| GTEx.Uterus | 70 | 566 | 0 |
| GTEx.Vagina | 79 | 635 | 0 |
| GTEx.Whole_Blood | 338 | 1991 | 0 |
| METSIM.ADIPOSE.RNASEQ | 563 | 4609 | 1 |
| YFS.BLOOD.RNAARR | 1264 | 4618 | 4 |
| CMC.BRAIN.RNASEQ | 452 | 5377 | 3 |
| NTR.BLOOD.RNAARR | 1247 | 2404 | 0 |

**Table S1: Tissue reference panels for predicting gene expression in TWAS.**

| Gene | Chr | TSS | TES | Best GWAS SNP | $Z_{GWAS}$ | Reference Tissue Panel | cis-$h_g^2$ | $P_{TWAS}$ |
|---|---|---|---|---|---|---|---|---|
| PJA2 | 5 | 108,670,409 | 108,745,675 | rs414724 | -6.4 | GTEx.Nerve_Tibial | 0.09 | 1.1E-07 |
| CTD-2587M2.1 | 5 | 108,572,821 | 108,662,070 | rs414724 | -6.4 | METSIM.ADIPOSE.RNASEQ | 0.18 | 3.3E-10 |
| MAN2A1 | 5 | 109,025,066 | 109,205,326 | rs414724 | -6.4 | GTEx.Nerve_Tibial | 0.07 | 2.4E-08 |
| SDK1 | 7 | 3,341,079 | 4,308,631 | rs10264803 | -6.0 | GTEx.Cells_Transformed_fibroblasts | 0.17 | 3.3E-09 |
| SDK1 | 7 | 3,341,079 | 4,308,631 | rs10264803 | -6.0 | GTEx.Esophagus_Muscularis | 0.08 | 6.4E-08 |
| EPDR1 | 7 | 37,960,162 | 37,991,542 | rs17171240 | 14.7 | GTEx.Lung | 0.15 | 6.4E-31 |
| EPDR1 | 7 | 37,960,162 | 37,991,542 | rs17171240 | 14.7 | GTEx.Adipose_Subcutaneous | 0.12 | 5.1E-23 |
| EPDR1 | 7 | 37,960,162 | 37,991,542 | rs17171240 | 14.7 | GTEx.Pancreas | 0.19 | 4.6E-18 |
| EPDR1 | 7 | 37,960,162 | 37,991,542 | rs17171240 | 14.7 | GTEx.Esophagus_Muscularis | 0.35 | 6.6E-14 |
| EPDR1 | 7 | 37,960,162 | 37,991,542 | rs17171240 | 14.7 | YFS.BLOOD.RNAARR | 0.18 | 1.4E-13 |
| EPDR1 | 7 | 37,960,162 | 37,991,542 | rs17171240 | 14.7 | GTEx.Nerve_Tibial | 0.30 | 5.0E-09 |
| EPDR1 | 7 | 37,960,162 | 37,991,542 | rs17171240 | 14.7 | GTEx.Artery_Tibial | 0.27 | 1.5E-08 |
| EPDR1 | 7 | 37,960,162 | 37,991,542 | rs17171240 | 14.7 | GTEx.Thyroid | 0.19 | 5.4E-08 |
| EPDR1 | 7 | 37,960,162 | 37,991,542 | rs17171240 | 14.7 | GTEx.Cells_Transformed_fibroblasts | 0.21 | 1.9E-07 |
| EPDR1 | 7 | 37,960,162 | 37,991,542 | rs17171240 | 14.7 | CMC.BRAIN.RNASEQ | 0.24 | 3.1E-07 |
| TRGC2 | 7 | 38,279,181 | 38,289,173 | rs17171240 | 14.7 | GTEx.Prostate | 0.37 | 1.4E-12 |
| SULF1 | 8 | 70,378,858 | 70,573,147 | rs542288 | 11.8 | GTEx.Artery_Aorta | 0.27 | 4.0E-25 |
| RSPO2 | 8 | 108,911,543 | 109,095,913 | rs612265 | -9.3 | CMC.BRAIN.RNASEQ | 0.12 | 1.2E-08 |
| EIF3E | 8 | 109,213,971 | 109,260,959 | rs612265 | -9.3 | CMC.BRAIN.RNASEQ | 0.07 | 7.6E-21 |
| EIF3E | 8 | 109,213,971 | 109,260,959 | rs612265 | -9.3 | YFS.BLOOD.RNAARR | 0.01 | 1.4E-15 |
| EIF3E | 8 | 109,213,971 | 109,260,959 | rs612265 | -9.3 | GTEx.Brain_Cerebellum | 0.16 | 1.8E-11 |
| EMC2 | 8 | 109,455,852 | 109,499,136 | rs612265 | -9.3 | GTEx.Muscle_Skeletal | 0.05 | 8.9E-12 |
| EMC2 | 8 | 109,455,852 | 109,499,136 | rs612265 | -9.3 | GTEx.Esophagus_Gastroesophageal_Junction | 0.09 | 1.3E-09 |
| EMC2 | 8 | 109,455,852 | 109,499,136 | rs612265 | -9.3 | GTEx.Brain_Cerebellum | 0.29 | 1.6E-08 |
| MRPL52 | 14 | 23,299,091 | 23,304,246 | rs1042704 | 7.3 | YFS.BLOOD.RNAARR | 0.44 | 1.1E-07 |
| NEDD4 | 15 | 56,119,116 | 56,285,944 | rs8032158 | 5.2 | GTEx.Artery_Tibial | 0.18 | 2.6E-07 |
| BCAR1 | 16 | 75,262,927 | 75,301,951 | rs977987 | 5.9 | GTEx.Artery_Aorta | 0.18 | 2.8E-07 |
| BCAR1 | 16 | 75,262,927 | 75,301,951 | rs977987 | 5.9 | GTEx.Esophagus_Muco | 0.16 | 3.6E-07 |

| | | | | | | | | |
|---|---|---|---|---|---|---|---|---|
| | | | | | | sa | | |
| CFDP1 | 16 | 75,327,607 | 75,467,387 | rs977987 | 5.9 | YFS.BLOOD.RNAARR | 0.21 | 5.6E-08 |
| TMEM170A | 16 | 75,480,922 | 75,498,584 | rs977987 | 5.9 | GTEx.Cells_EBV-transformed_lymphocytes | 0.13 | 1.4E-09 |
| TMEM170A | 16 | 75,480,922 | 75,498,584 | rs977987 | 5.9 | GTEx.Skin_Sun_Exposed_Lower_leg | 0.14 | 2.7E-08 |
| TMEM170A | 16 | 75,480,922 | 75,498,584 | rs977987 | 5.9 | GTEx.Skin_Not_Sun_Exposed_Suprapubic | 0.09 | 3.9E-08 |
| TMEM106A* | 17 | 41,363,845 | 41,372,057 | rs4793248 | 4.1 | GTEx.Breast_Mammary_Tissue | 0.12 | 1.2E-07 |
| ATXN10 | 22 | 46,067,677 | 46,241,187 | rs34088184 | 13.8 | GTEx.Cells_Transformed_fibroblasts | 0.17 | 1.7E-07 |
| LINC00899 | 22 | 46,435,786 | 46,440,748 | rs34088184 | 13.8 | GTEx.Adipose_Subcutaneous | 0.17 | 2.8E-32 |
| LINC00899 | 22 | 46,435,786 | 46,440,748 | rs34088184 | 13.8 | GTEx.Muscle_Skeletal | 0.24 | 1.6E-26 |
| LINC00899 | 22 | 46,435,786 | 46,440,748 | rs34088184 | 13.8 | GTEx.Cells_Transformed_fibroblasts | 0.18 | 1.9E-26 |
| LINC00899 | 22 | 46,435,786 | 46,440,748 | rs34088184 | 13.8 | GTEx.Artery_Tibial | 0.30 | 5.6E-23 |
| LINC00899 | 22 | 46,435,786 | 46,440,748 | rs34088184 | 13.8 | GTEx.Esophagus_Muscularis | 0.36 | 2.3E-15 |
| LINC00899 | 22 | 46,435,786 | 46,440,748 | rs34088184 | 13.8 | GTEx.Lung | 0.21 | 5.7E-15 |
| LINC00899 | 22 | 46,435,786 | 46,440,748 | rs34088184 | 13.8 | GTEx.Adrenal_Gland | 0.68 | 9.2E-15 |
| LINC00899 | 22 | 46,435,786 | 46,440,748 | rs34088184 | 13.8 | GTEx.Artery_Coronary | 0.56 | 1.6E-09 |
| LINC00899 | 22 | 46,435,786 | 46,440,748 | rs34088184 | 13.8 | GTEx.Nerve_Tibial | 0.35 | 3.4E-08 |

**Table S2: Significant tissue-specific gene expression models from TWAS.** These are the 43 significant ($P_{TWAS} \leq \frac{0.05}{98147}$) tissue-specific gene models representing 18 unique genes. The novel risk region is from the *TMEM106A* model significant in the GTEx breast mammary tissue, indicated by "*".

| Reference Tissue Panel | cis-$h_g^2$ | Best GWAS SNP | $Z_{GWAS}$ | $Z_{TWAS}$ | $P_{TWAS}$ |
|---|---|---|---|---|---|
| GTEx.Artery_Aorta | 0.24 | rs4793248 | 4.12 | 3.42 | 6.2E-04 |
| GTEx.Artery_Coronary | 0.17 | rs4793248 | 4.12 | 4.16 | 3.1E-05 |
| GTEx.Artery_Tibial | 0.16 | rs4793248 | 4.12 | 3.37 | 7.4E-04 |
| GTEx.Breast_Mammary_Tissue | 0.12 | rs4793248 | 4.12 | 5.3 | 1.2E-07 |
| GTEx.Cells_EBV-transformed_lymphocytes | 0.46 | rs4793248 | 4.12 | -3.12 | 0.0018 |
| GTEx.Heart_Atrial_Appendage | 0.08 | rs4793248 | 4.12 | 4.8 | 1.6E-06 |
| GTEx.Nerve_Tibial | 0.15 | rs4793248 | 4.12 | 3.63 | 2.8E-04 |
| GTEx.Pituitary | 0.15 | rs4793248 | 4.12 | 3.76 | 1.7E-04 |
| GTEx.Skin_Not_Sun_Exposed_Suprapubic | 0.07 | rs4793248 | 4.12 | 1.9 | 0.0571 |
| GTEx.Small_Intestine_Terminal_Ileum | 0.15 | rs4793248 | 4.12 | 4.29 | 1.8E-05 |
| GTEx.Vagina | 0.41 | rs4793248 | 4.12 | 1.73 | 0.0845 |
| YFS.BLOOD.RNAARR | 0.02 | rs4793248 | 4.12 | -1.3 | 0.1948 |

**Table S3: All tissue-specific gene models for the *TMEM106A*.** Here we have the 12 different tissues that *TMEM106A* (chr17:41363845-41372057) expression was tested with for association with DD. Only the GTEx breast mammary tissue model was significant.

| Trait | LDSC $\hat{r}_g$ | s.e. | ρ-HESS $\hat{r}_g$ | s.e. |
|---|---|---|---|---|
| BMI | -0.196 | 0.04 | -0.229 | 0.02 |
| HDL | 0.133 | 0.04 | 0.093 | 0.03 |
| TG | -0.139 | 0.04 | -0.109 | 0.03 |
| T2D | -0.182 | 0.05 | -0.112 | 0.04 |

**Table S4: Genetic correlation for four traits with DD using LDSC and ρ-HESS.** Here are the genome-wide genetic correlation estimates from LDSC and ρ-HESS for four traits. Pearson's correlation between these estimates is 0.94.

| Tissue | # Genes | Beta | s.e. | $P_{GP,T}$ |
|---|---|---|---|---|
| Cells_Transformed_fibroblasts | 16517 | 0.025 | 0.008 | 0.001 |
| Nerve_Tibial | 16517 | 0.034 | 0.012 | 0.002 |
| Adipose_Subcutaneous | 16517 | 0.029 | 0.012 | 0.009 |
| Breast_Mammary_Tissue | 16517 | 0.033 | 0.014 | 0.012 |
| Cervix_Ectocervix | 16517 | 0.03 | 0.015 | 0.021 |
| Artery_Tibial | 16517 | 0.023 | 0.012 | 0.024 |
| Artery_Aorta | 16517 | 0.023 | 0.012 | 0.025 |
| Vagina | 16517 | 0.021 | 0.013 | 0.048 |
| Small_Intestine_Terminal_Ileum | 16517 | 0.015 | 0.009 | 0.055 |
| Uterus | 16517 | 0.018 | 0.013 | 0.078 |
| Minor_Salivary_Gland | 16517 | 0.016 | 0.011 | 0.08 |
| Cervix_Endocervix | 16517 | 0.019 | 0.014 | 0.081 |
| Adipose_Visceral_Omentum | 16517 | 0.018 | 0.013 | 0.081 |
| Skin_Sun_Exposed_Lower_leg | 16517 | 0.012 | 0.009 | 0.083 |
| Artery_Coronary | 16517 | 0.018 | 0.014 | 0.088 |
| Esophagus_Mucosa | 16517 | 0.011 | 0.008 | 0.095 |
| Fallopian_Tube | 16517 | 0.018 | 0.014 | 0.097 |
| Skin_Not_Sun_Exposed_Suprapubic | 16517 | 0.011 | 0.009 | 0.101 |
| Thyroid | 16517 | 0.014 | 0.011 | 0.109 |
| Whole_Blood | 16517 | 0.008 | 0.006 | 0.113 |
| Prostate | 16517 | 0.016 | 0.013 | 0.115 |
| Cells_EBV-transformed_lymphocytes | 16517 | 0.006 | 0.006 | 0.14 |
| Spleen | 16517 | 0.008 | 0.008 | 0.148 |
| Colon_Transverse | 16517 | 0.012 | 0.012 | 0.154 |
| Lung | 16517 | 0.01 | 0.011 | 0.164 |
| Muscle_Skeletal | 16517 | 0.006 | 0.008 | 0.232 |
| Bladder | 16517 | 0.009 | 0.014 | 0.252 |
| Esophagus_Gastroesophageal_Junction | 16517 | 0.008 | 0.015 | 0.286 |
| Colon_Sigmoid | 16517 | 0.008 | 0.014 | 0.292 |
| Ovary | 16517 | 0.004 | 0.011 | 0.368 |
| Esophagus_Muscularis | 16517 | 0.004 | 0.014 | 0.393 |
| Kidney_Cortex | 16517 | -0.002 | 0.011 | 0.592 |
| Testis | 16517 | -0.002 | 0.006 | 0.603 |
| Pancreas | 16517 | -0.006 | 0.01 | 0.725 |
| Brain_Spinal_cord_cervical_c-1 | 16517 | -0.006 | 0.01 | 0.731 |
| Stomach | 16517 | -0.008 | 0.014 | 0.736 |
| Brain_Cerebellum | 16517 | -0.005 | 0.007 | 0.767 |

| | | | | |
|---|---|---|---|---|
| Adrenal_Gland | 16517 | -0.008 | 0.011 | 0.784 |
| Brain_Cerebellar_Hemisphere | 16517 | -0.006 | 0.007 | 0.804 |
| Liver | 16517 | -0.006 | 0.007 | 0.821 |
| Brain_Hippocampus | 16517 | -0.018 | 0.009 | 0.976 |
| Heart_Atrial_Appendage | 16517 | -0.021 | 0.011 | 0.978 |
| Brain_Substantia_nigra | 16517 | -0.02 | 0.01 | 0.98 |
| Brain_Amygdala | 16517 | -0.019 | 0.009 | 0.982 |
| Brain_Cortex | 16517 | -0.017 | 0.008 | 0.985 |
| Heart_Left_Ventricle | 16517 | -0.022 | 0.01 | 0.987 |
| Brain_Anterior_cingulate_cortex_BA24 | 16517 | -0.019 | 0.008 | 0.989 |
| Brain_Frontal_Cortex_BA9 | 16517 | -0.018 | 0.008 | 0.991 |
| Brain_Hypothalamus | 16517 | -0.021 | 0.009 | 0.991 |
| Brain_Caudate_basal_ganglia | 16517 | -0.021 | 0.009 | 0.992 |
| Brain_Putamen_basal_ganglia | 16517 | -0.022 | 0.009 | 0.992 |
| Brain_Nucleus_accumbens_basal_ganglia | 16517 | -0.021 | 0.008 | 0.993 |
| Pituitary | 16517 | -0.029 | 0.01 | 0.999 |

**Table S7: FUMA gene property analysis for 53 GTEx tissues.**
